# Coffee remodels the global histone acetylation landscape with reduced H3K27ac at the *MYC* promoter

**DOI:** 10.64898/2026.08.01.742200

**Authors:** Yangsu Cho, Makiko Miyota-Yui, Yaeko Nakajima-Takagi, Takako Yokomizo, Motohiko Oshima, Atsushi Iwama, Kazumasa Aoyama

## Abstract

Coffee consumption has been associated with reduced risks of multiple chronic diseases, including cancer; however, whether coffee regulates the epigenetic landscape remains largely unknown. Here, we show that coffee extensively remodels the histone acetylation landscape in human cells. Quantitative histone modification proteomics revealed a global reduction in active histone acetylation marks following coffee treatment. Mechanistically, coffee reduced the abundance of the histone acetyltransferases GCN5 and MOF, decreased 14-3-3 proteins, promoted the nuclear accumulation of class II histone deacetylases, and induced histone hypoacetylation that was reversed by the histone deacetylase inhibitor trichostatin A. Genome-wide H3K27ac profiling revealed widespread loss of promoter and enhancer acetylation, accompanied by broad transcriptional reprogramming. Integrative analyses of ChIP-seq, RNA-seq, proteomics and biochemical validation consistently demonstrated suppression of the MYC transcriptional program, including reduced *MYC* expression, decreased H3K27ac at MYC target promoters, downregulation of canonical MYC target genes and attenuation of MYC target gene signatures. Functionally, coffee inhibited cancer cell proliferation with minimal cell death. Collectively, our findings identify whole coffee as a previously unrecognized epigenetic modulator that remodels the histone acetylation landscape in association with suppression of the MYC transcriptional program, providing a molecular framework for understanding the epigenetic basis underlying the anti-tumor effects of coffee.

## Introduction

Coffee is one of the most widely consumed beverages worldwide and has consistently been associated with reduced risks of multiple chronic diseases^1,2^, including cardiovascular disease, type 2 diabetes, neurodegenerative disorders, and several types of cancer ^3,4^. Epidemiological studies and meta-analyses have suggested that habitual coffee consumption is associated with decreased cancer incidence and mortality, although the molecular mechanisms underlying these protective effects remain incompletely understood.

Coffee contains hundreds of bioactive compounds, including caffeine, chlorogenic acids, trigonelline, diterpenes, and various phenolic metabolites. These compounds have been reported to exert antioxidant, anti-inflammatory, and metabolic effects through modulation of multiple signaling pathways ^5,6^. However, because whole coffee represents a complex mixture of numerous biologically active molecules, it remains unclear whether coffee also regulates chromatin structure and epigenetic states at the genome-wide level.

Epigenetic regulation, including histone acetylation and methylation, plays central roles in controlling gene expression, cell identity, proliferation, and tumorigenesis ^7–10^. Among active histone modifications, H3K27ac serves as a hallmark of active promoters and enhancers, whereas H3K18ac and H3K36ac are also closely associated with transcriptionally active chromatin ^11,12^. Histone acetylation is dynamically regulated by the balance between histone acetyltransferases (HATs) and histone deacetylases (HDACs), and disruption of this balance contributes to cancer development and many other human diseases ^13–16^.

Several studies have suggested that dietary components can influence epigenetic regulation^17^, and DNA methylation changes associated with coffee consumption have been reported in human epigenome-wide association studies ^18,19^. A recent comprehensive review further summarized the epigenetic effects of coffee and its major constituents, including caffeine, chlorogenic acid, and caffeic acid, concluding that current evidence primarily supports effects on DNA methylation, non-coding RNAs, and selected histone modifications induced by individual coffee-derived compounds ^20^. Genome-wide analyses have shown that individual coffee-derived compounds, including caffeine and chlorogenic acid, can modulate H3K27 acetylation in specific biological contexts ^21,22^. Nevertheless, the epigenetic effects of whole coffee remain poorly understood. To the best of our knowledge, no study has comprehensively investigated how whole coffee remodels the histone acetylation landscape at the genome-wide level by integrating quantitative histone modification proteomics, chromatin profiling, transcriptomics, and proteomics.

In the present study, we combined quantitative histone modification proteomics, quantitative proteomics, ChIP-seq, RNA-seq, biochemical analyses, and functional assays to investigate the effects of coffee on chromatin regulation. We demonstrate that coffee induces a global reduction in active histone marks, including H3K27ac, H3K18ac, H3K36ac, and H3K4me3. Coffee-induced histone hypoacetylation is mediated through coordinated regulation of histone acetyltransferases and histone deacetylases, resulting in widespread loss of promoter-associated H3K27ac, suppression of the MYC transcriptional program, and reduced cancer cell proliferation. These findings identify coffee as a previously unrecognized epigenetic modulator and provide a molecular framework linking coffee consumption to chromatin remodeling and transcriptional regulation.

## Materials and Methods

### Cell culture

K562 cells were cultured in RPMI 1640 medium supplemented with 10% fetal bovine serum (FBS), 100 U/mL penicillin, and 100 μg/mL streptomycin. TIG-1 human diploid fibroblasts (JCRB0501) were obtained from the Japanese Collection of Research Bioresources (JCRB) Cell Bank (Osaka, Japan) and cultured in low-glucose DMEM (1.0 g/L glucose) supplemented with 10% fetal bovine serum (FBS), 100 U/mL penicillin, and 100 μg/mL streptomycin. MCF-7, A549, A431, and HCT116 cells were kindly provided by Professor Noritaka Yamaguchi (Meiji Pharmaceutical University, Tokyo, Japan) and cultured in high-glucose DMEM (4.5 g/L glucose) supplemented with 10% fetal bovine serum (FBS), 100 U/mL penicillin, and 100 μg/mL streptomycin. All cells were maintained at 37°C in a humidified incubator with 5% CO₂.

### Coffee preparation

Six commercially available coffee products were used in this study: Starbucks House Blend, Starbucks Decaf House Blend (Starbucks Coffee Company, Seattle, WA, USA), TULLY’S BARISTA’S BLACK (Ito En, Tokyo, Japan), UCC BLACK Unsweetened (UCC Ueshima Coffee Co., Ltd., Kobe, Japan), GEORGIA THE PREMIUM (The Coca-Cola Company, Tokyo, Japan), and BOSS Black (Suntory Beverage & Food Ltd., Tokyo, Japan). Unless otherwise indicated, Starbucks House Blend was used for subsequent experiments, and Starbucks Decaf House Blend was used for decaffeinated coffee experiments. Starbucks House Blend and Starbucks Decaf House Blend were prepared according to the manufacturer’s instructions by brewing 10 g of ground coffee with 180 mL of freshly boiled water using a commercially available paper coffee filter. After cooling to room temperature, the coffee extracts were sterilized by filtration through a 0.22-μm membrane filter, aliquoted, and stored at −80°C until use. Ready-to-drink coffee products (TULLY’S BARISTA’S BLACK, UCC BLACK Unsweetened, GEORGIA THE PREMIUM, and BOSS Black) were aseptically aliquoted into sterile tubes and stored at −80°C until use. Coffee extracts were added to the culture medium at a final concentration of 5% (v/v). This concentration was selected based on previous cell culture studies investigating the biological effects of coffee^23–25^.

### Estimation of coffee-derived compound concentrations

The concentrations of major coffee-derived compounds in brewed coffee were estimated from previously published compositional analyses. The concentration of total chlorogenic acids was based on the average total chlorogenic acid content (mg/g ground coffee) of regular brewed coffees reported by Fujioka and Shibamoto, who prepared brewed coffee from 12.5 g of ground coffee and 450 mL of water^26^. The concentrations of caffeine, trigonelline, caffeic acid, *p*-coumaric acid, nicotinic acid, 5-hydroxymethylfurfural (5-HMF), and theobromine were estimated by multiplying the total chlorogenic acid content by the corresponding compound-to-total chlorogenic acid ratios reported by Rodrigues and Bragagnolo^27^. Estimated amounts (mg/g ground coffee) were converted to concentrations in our coffee extract, which was prepared by brewing 10 g of ground coffee with 180 mL of hot water using a paper filter, and expressed as mM using the molecular weight of each compound. The concentration of pyrocatechol was estimated from the value reported by Funakoshi-Tago et al. ^23^ and corrected according to the coffee powder-to-water ratio to match the brewing conditions used in the present study. The average values were used to determine the concentrations of individual coffee-derived compounds used in the experiments, whereas the reported minimum and maximum chlorogenic acid contents were used to estimate the concentration ranges shown in Fig. 3E and Supplementary Table S1. The detailed calculations are provided in Supplementary Table S1, and the resulting estimated concentrations together with the experimental concentrations are summarized in Fig. 3E.

### Antibodies

The following antibodies were used: anti-H3K27ac (#8173, Cell Signaling Technology), anti-H3K18ac (#13998, Cell Signaling Technology), anti-H3K36ac (#27683, Cell Signaling Technology), anti-H3K4me3 (#8580, Abcam), anti-H3K4me1 (#8895, Cell Signaling Technology), anti-MOF (MYST1) (#46862, Cell Signaling Technology), anti-GCN5 (GCN5L2) (#3305, Cell Signaling Technology), anti-HAT1 (#41490, Cell Signaling Technology), anti-Phospho-HDAC Class II (#3443, Cell Signaling Technology), anti-14-3-3 (pan) (#9542, Cell Signaling Technology), anti-HDAC3 (#3949, Cell Signaling Technology), anti-HDAC4 (#7628, Cell Signaling Technology), anti-Caspase-3 (#9662, Cell Signaling Technology), anti-Cleaved Caspase-3 (#9661, Cell Signaling Technology), anti-Mouse IgG-HRP (light chain specific, #115-035-174, Jackson ImmunoResearch), and anti-Rabbit IgG-HRP (light chain specific, #211-032-171, Jackson ImmunoResearch), anti-Rabbit IgG-Alexa Fluor 488 (#AB150077, Abcam) antibodies.

### Western blotting

Western blot analysis was carried out essentially as described previously^28,29^. Cells were washed with PBS and lysed in SDS lysis buffer (2% SDS, 20 mM Tris-HCl, pH 8.0) ^30^. Cell lysates were sonicated, mixed with an equal volume of 2× SDS sample buffer, and heated at 95°C for 10 min. Proteins were separated by SDS-PAGE and transferred onto PVDF membranes (PALL). After blocking with 1% BSA, membranes were incubated with the indicated primary and secondary antibodies. Signals were visualized using Immobilon Western chemiluminescence reagent (Millipore) and acquired with a ChemiDoc™ Touch imaging system (Bio-Rad). For sequential reprobing, antibodies were removed by incubation in 0.2 M glycine-HCl buffer (pH 2.5), and/or HRP activity was quenched using 0.1% NaN3, as described previously ^31,32^. Image processing for figure preparation was performed using GIMP software, and densitometric analysis of band intensities was conducted using ImageJ software ^33^.

### Subcellular fractionation

Cytoplasmic and nuclear fractions were prepared using a hypotonic lysis method as previously described^31,34^ with minor modifications. Briefly, cells were washed twice with ice-cold PBS and resuspended in hypotonic lysis buffer (20 mM Tris-HCl, pH 8.0, 10 mM NaCl, 3 mM MgCl₂, Complete EDTA-free Protease Inhibitor Cocktail (Roche), and PhosSTOP phosphatase inhibitor (Roche)). After incubation on ice for 5 min, NP-40 was added to a final concentration of 0.1%, and the cell suspension was gently mixed and centrifuged at 5,000 × g for 5 min at 4°C. The resulting supernatant was collected as the cytoplasmic fraction. The nuclear pellet was washed twice with hypotonic buffer containing 0.1% NP-40 and lysed in 2% SDS lysis buffer (20 mM Tris-HCl, pH 8.0, 2% SDS), followed by brief sonication to reduce viscosity. Equal volumes of 2× SDS sample buffer were added to both cytoplasmic and nuclear fractions, and samples were heated at 95°C for 10 min. The fractions were subsequently analyzed by western blotting.

### Immunofluorescence microscopy

Immunofluorescence analysis was carried out largely according to previously described procedures^35,36^ with minor modifications. For K562 cells, cells were collected into 1.5-mL microcentrifuge tubes, and all fixation and staining procedures were performed in suspension. Following staining, cells were mixed with Fluoro-KEEPER Antifade Reagent, Non-Hardening Type with DAPI (Nacalai Tesque), placed onto glass coverslips (22 × 22 mm, Matsunami), mounted with microscope slides, and sealed with nail polish. For TIG1 cells, sterile glass coverslips (22 × 22 mm, Matsunami) were placed in 6-well plates, and cells were seeded directly onto the coverslips. All fixation and staining procedures were subsequently performed on the coverslips. After staining, the coverslips were mounted onto microscope slides using Fluoro-KEEPER Antifade Reagent, Non-Hardening Type with DAPI (Nacalai Tesque) and sealed with nail polish.

Both K562 and TIG1 cells were fixed with 4% paraformaldehyde for 20 min at room temperature and washed three times with PBS. Cells were permeabilized and blocked with PBS containing 0.5% Triton X-100 and 1% BSA for 20 min. Samples were incubated for 1h at room temperature or overnight at 4°C with anti-H3K27ac antibody (#8173, Cell Signaling Technology) diluted in blocking buffer. After washing three times with PBS, anti-Rabbit IgG-Alexa Fluor 488 (#AB150077, Abcam) was applied for 30–60 min at room temperature in the dark. Fluorescence images were acquired using an FV4000 confocal laser scanning microscope (Evident, Japan).

Quantification of H3K27ac fluorescence intensity was performed using ImageJ software (NIH) as described previously^37,38^. Nuclear regions were defined based on DAPI staining, and the mean fluorescence intensity of H3K27ac within each nucleus was measured after background subtraction.

### Proteomic analyses

For both histone modification profiling and global proteome analysis, protein samples were subjected to liquid chromatography–tandem mass spectrometry (LC–MS/MS)-based proteomic analysis by commercial service providers.

Histone post-translational modifications were analyzed using the Mod Spec platform (Active Motif, Carlsbad, CA, USA). Histones were acid-extracted, derivatized, digested, and analyzed by LC–MS/MS as previously described by the service provider. Relative abundance of each histone peptide modification was quantified as the percentage of the total signal corresponding to the respective peptide. One biological sample was analyzed with three technical LC–MS/MS replicates.

For global protein expression analysis, whole-cell lysates were subjected to LC–MS/MS analysis by Kazusa DNA Research Institute (Kisarazu, Japan). Protein identification and label-free quantification were performed using the institute’s standard proteomic workflow.

### Chromatin immunoprecipitation sequence (ChIP-seq)

Chromatin immunoprecipitation (ChIP) was performed as described previously ^39,40^ with minor modifications. Briefly, cells were fixed with 0.5% formaldehyde, and the reaction was quenched with glycine. Cell pellets were lysed in ChIP lysis buffer (10 mM Tris-HCl, pH 8.0, 200 mM NaCl, 1 mM CaCl₂, and 0.5% NP-40, supplemented with Complete Mini EDTA-free protease inhibitor cocktail (Roche)). The lysates were briefly sonicated to disrupt the cells and nuclei, followed by digestion with micrococcal nuclease (MNase; New England Biolabs) at a final concentration of 2,000 gel units/mL for 40 min at 37°C. The reaction was terminated by the addition of 10 mM EDTA, followed by the addition of RIPA lysis buffer (50 mM Tris-HCl, pH 8.0, 150 mM NaCl, 2 mM EDTA, 1% NP-40, 0.5% sodium deoxycholate, and 0.1% SDS). The lysates were further sonicated to release MNase-digested chromatin from the nuclei and solubilize chromatin. After centrifugation, the soluble chromatin fraction was collected, and an aliquot was reserved as the input control. Dynabeads M-280 Sheep anti-Rabbit IgG (Thermo Fisher Scientific) were pre-incubated with anti-H3K27ac antibody (#8173, Cell Signaling Technology) and incubated with the chromatin overnight at 4°C. The beads were washed sequentially with ChIP wash buffer (10 mM Tris-HCl, pH 8.0, 500 mM NaCl, 1 mM CaCl₂, and 0.5% NP-40) followed by TE buffer (10 mM Tris-HCl, pH 8.0, and 1 mM EDTA). Immunoprecipitated chromatin was eluted using elution buffer (50 mM Tris-HCl, pH 8.0, 10 mM EDTA, and 1% SDS). Cross-links were reversed by incubation at 65°C, followed by RNase A and Proteinase K treatment. DNA was purified using the MinElute PCR Purification Kit (Qiagen). ChIP-seq libraries were prepared using the ThruPLEX DNA-Seq Kit (Takara Bio) according to the manufacturer’s instructions. Following adaptor ligation, libraries were size-selected using AMPure XP beads and sequenced on an Illumina platform.

To account for the global reduction in H3K27ac induced by coffee treatment, H3K27ac ChIP-seq signals were normalized using the relative global H3K27ac level, as previously described ^39,41^. In the present study, the relative global H3K27ac level was estimated from the amount of H3K27ac-immunoprecipitated DNA (Fig. 6A), and normalized ChIP-seq signal intensities were multiplied by this correction factor.

For Fig. 5, publicly available ChIP-seq datasets were obtained from ChIP-Atlas^42^ and included H3K27ac (SRX8848926), H3K18ac (SRX3431313), and H3K4me3 (ERX989282). These datasets were analyzed using the same bioinformatics pipeline as our ChIP-seq data.

### RNA sequence (RNA-seq)

Cell lysates in Sepasol-RNA I Super G (Nacalai Tesque) were submitted to the Kazusa DNA Research Institute for RNA extraction, RNA quality assessment, library preparation, sequencing, and gene expression quantification. RNA integrity was evaluated using an Agilent 2100 Bioanalyzer (Agilent Technologies). Sequencing libraries were prepared from 500 ng of total RNA using the QuantSeq 3′ mRNA-Seq Library Prep Kit V2 (FWD) with UDI (Lexogen). Single-end sequencing (75 bp) was performed on an Illumina NextSeq 500 platform. Gene expression levels were quantified as reads per million (RPM) by the Kazusa DNA Research Institute.

### Gene set enrichment analysis (GSEA)

Gene set enrichment analysis (GSEA) was performed using the Broad Institute GSEA software, as described previously ^43^. Differentially expressed genes identified from RNA-seq analysis were ranked according to expression changes between control and coffee-treated cells. Hallmark gene sets from the Molecular Signatures Database (MSigDB) were used for pathway enrichment analysis^44^.

### Gene Ontology (GO) Enrichment Analysis

Functional enrichment analysis was conducted using the Database for Annotation, Visualization, and Integrated Discovery (DAVID) platform ^45,46^, as described previously ^36,39^ with minor modifications. Lists of proteins or differentially expressed genes were analyzed using official gene symbols, and enrichment was evaluated within the Gene Ontology (GO) Biological Process category under default human background settings.

### Reverse transcription quantitative PCR (RT–qPCR)

Total RNA was extracted using Sepasol-RNA I Super G (Nacalai Tesque, Kyoto, Japan). cDNA was synthesized from 1 μg of total RNA using RT Ace (Toyobo, Osaka, Japan). Quantitative PCR (qPCR) was performed using Luna® Universal qPCR Master Mix (New England Biolabs, Ipswich, MA, USA). Relative mRNA expression was normalized to ACTB using the ΔΔCt method. The primer sequences were as follows: MYC (forward, 5′-CAGGACCCGCTTCTCTGAAA-3′; reverse, 5′-CTAACGTTGAGGGGCATCGT-3′), ODC1 (forward, 5′-CGCCTGGGCGCTCTGA-3′; reverse, 5′-TTTCCAGCTTCTCACAAAGGC-3′), LDHA (forward, 5′-CGCCGATTCCGGATCTCATT-3′; reverse, 5′-AGCTGATCCTTTTAGAGTTGCCA-3′), MCM7 (forward, 5′-CCCCTCCCAGTTTGAACCTC-3′; reverse, 5′-TCTCGCCTCATCTCCACGTA-3′), PRPS2 (forward, 5′-CAATGCCTGCAAAGATTGCGT-3′; reverse, 5′-TTTGCAGAAATTGGGGCACG-3′), and ACTB (forward, 5′-AAGTTCACAATGTGGCCGAG-3′; reverse, 5′-AGTGGGGTGGCTTTTAGGATG-3′).

### Cell proliferation and viability assays

Cell proliferation was evaluated by direct cell counting and WST assays, as described previously ^47,48^ with minor modifications. K562 or TIG-1 cells were seeded at the indicated densities and treated with coffee for the indicated periods. Cell numbers were determined using a hemocytometer. Cell metabolic activity was assessed using Cell Count Reagent SF (#07553-86, Nacalai Tesque, Kyoto, Japan) according to the manufacturer’s instructions. Absorbance was measured at 450 nm using a microplate reader. Cell viability was evaluated by the trypan blue exclusion assay. Following treatment, cells were mixed with 0.4% trypan blue solution, and viable and non-viable cells were counted using a hemocytometer. Cell viability was expressed as the percentage of trypan blue-negative cells.

### Deposition of data

The mass spectrometry proteomics data have been deposited to the jPOST repository with the dataset identifier JPST004748. RNA-seq and ChIP-seq data have been deposited in the NCBI Gene Expression Omnibus (GEO) under accession number GSE337644 (linked SRA BioProject: PRJNA1464642).

## Results

### Coffee globally reduces histone acetylation in human cells

To determine whether coffee alters the epigenetic landscape, we first performed quantitative histone modification proteomic analysis in K562 cells treated with 5% (v/v) coffee for 24 h, using a concentration previously employed in cell culture studies^23–25^ (Fig. 1A). Comprehensive profiling identified widespread alterations in histone modifications following coffee treatment (Fig. 1B, C). Among the detected modifications, histone acetylation marks were predominantly decreased, whereas relatively few histone methylation marks were affected (Fig. 1B). Heatmap analysis further demonstrated a global reduction in histone acetylation, while most histone methylation marks remained largely unchanged (Fig. 1C), suggesting that coffee primarily induces histone hypoacetylation rather than a global shift from histone acetylation to histone methylation. Quantification of the most strongly altered modifications revealed that multiple lysine acetylation sites, including H3K18ac, H3K27ac, and H3K36ac, were among the most prominently reduced histone modifications following coffee treatment (Fig. 1D).

**Figure 1.**
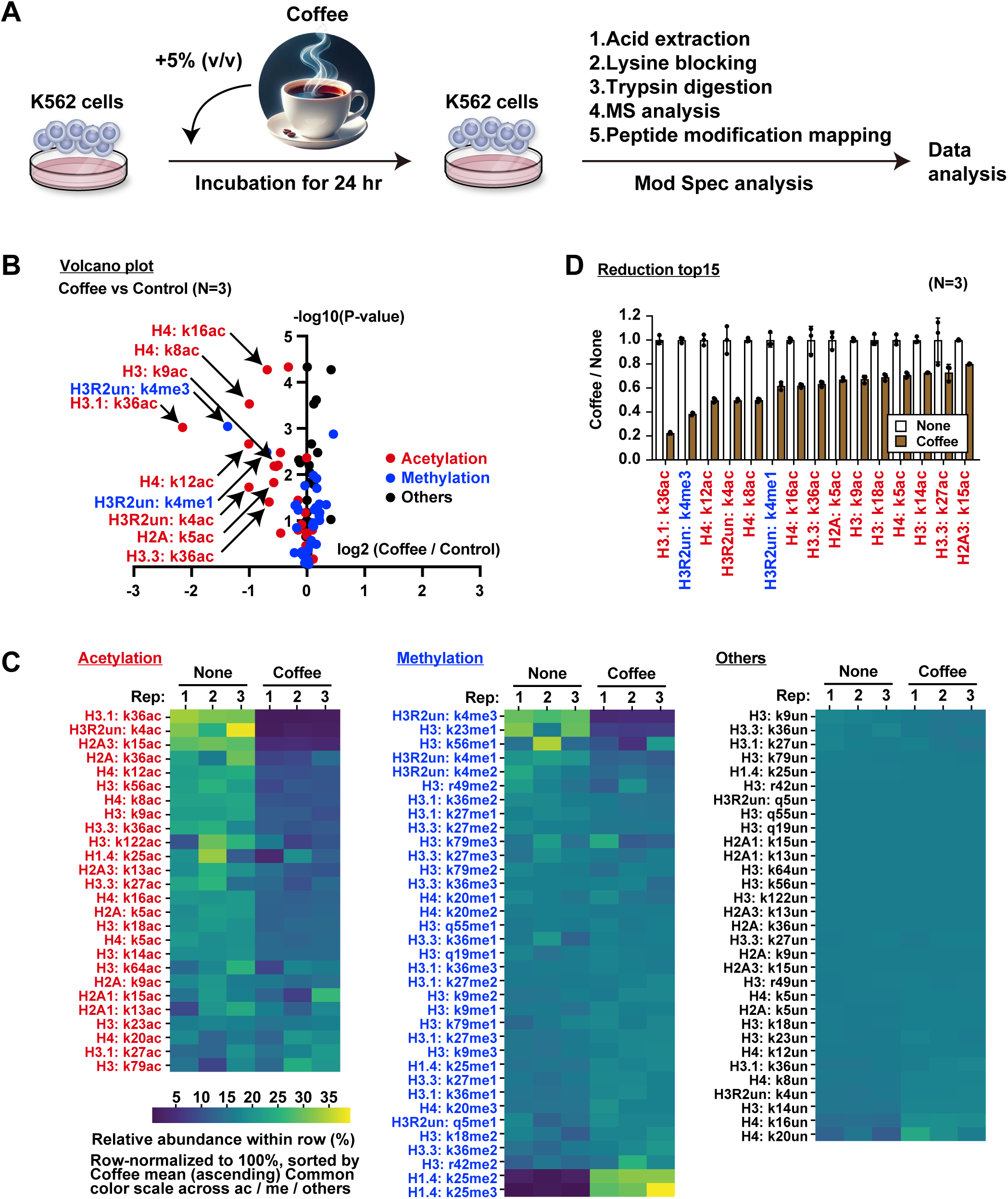
Global remodeling of the histone modification landscape induced by coffee revealed by quantitative histone modification proteomics. (A) Schematic overview of the experimental workflow. K562 cells were treated with 5% (v/v) coffee for 24 h. Histones were acid-extracted, chemically derivatized, digested with trypsin, and subjected to quantitative histone modification proteomic analysis by mass spectrometry. (B) Volcano plot showing changes in histone modifications following coffee treatment in K562 cells (n = 3). Histone acetylation, histone methylation, and other histone modifications are indicated in red, blue, and black, respectively. Representative significantly altered histone modifications are labeled. (C) Heat maps showing the relative abundance of histone acetylation, histone methylation, and other histone modifications in individual technical replicates. Values were row-normalized to 100% and ordered according to the mean abundance in coffee-treated samples. A common color scale was applied within each modification category. In the heatmaps, “un” indicates the unmodified form of the corresponding histone residue. (D) Relative abundance of the top 15 histone modifications exhibiting the greatest reduction following coffee treatment. Data are presented as mean ± SD (n = 3 biological replicates).

To validate these proteomic findings, we examined representative histone modifications by immunoblotting. In K562 cells, coffee treatment significantly reduced the levels of H3K18ac, H3K27ac, and H3K36ac, whereas H3K4me1 was largely unaffected (Fig. 2A). H3K4me3 also showed a moderate reduction, consistent with the proteomic analysis (Fig. 2A). In contrast, coffee did not significantly alter these histone modifications in the normal human fibroblast cell line TIG-1 (Fig. 2B), indicating that the effect of coffee on histone acetylation is cell type-dependent. Consistent with these observations, coffee also reduced H3K27ac levels in multiple human cancer cell lines, including MCF7, A549, and A431 cells, whereas no significant change was observed in HCT116 cells (Fig. 2C). Immunofluorescence analysis further confirmed a marked reduction in nuclear H3K27ac staining in K562 cells but not in TIG-1 cells following coffee treatment (Fig. 2D). Collectively, these findings demonstrate that coffee preferentially induces global histone hypoacetylation in multiple human cancer cell types, establishing histone hypoacetylation as a prominent epigenetic response to coffee treatment.

**Figure 2.**
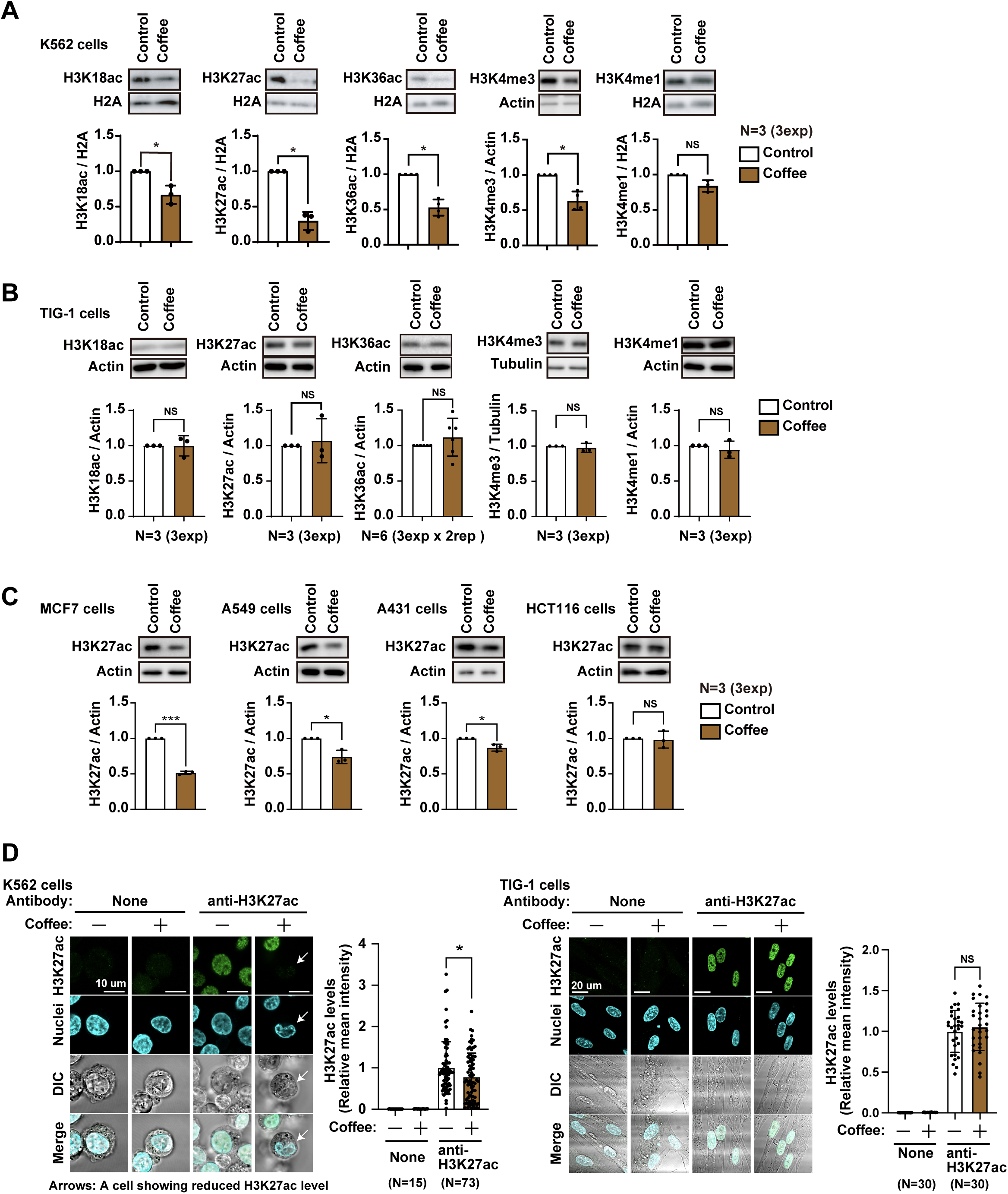
Cell type-dependent regulation of histone acetylation by coffee. **(A)** Immunoblot analysis of H3K18ac, H3K27ac, H3K36ac, H3K4me3, and H3K4me1 in K562 cells treated with coffee for 24 h. Representative immunoblots and corresponding quantification are shown. Data are presented as mean ± SD from three independent experiments. **(B)** Immunoblot analysis of H3K18ac, H3K27ac, H3K36ac, H3K4me3, and H3K4me1 in TIG-1 cells following coffee treatment. Representative immunoblots and corresponding quantification are shown. Data are presented as mean ± SD from three independent experiments unless otherwise indicated. **(C)** Immunoblot analysis of H3K27ac in MCF7, A549, A431, and HCT116 cells following coffee treatment. Representative immunoblots and corresponding quantification are shown. Data are presented as mean ± SD from three independent experiments. **(D)** Immunofluorescence staining of H3K27ac in K562 and TIG-1 cells following coffee treatment. Representative images and quantification of nuclear H3K27ac fluorescence intensity are shown. Arrowheads indicate representative cells with reduced nuclear H3K27ac staining. Scale bars, 10 μm (K562) and 20 μm (TIG-1).

### Multiple coffee preparations induce histone hypoacetylation, whereas individual coffee-derived components only partially reproduce this effect

To determine whether the observed histone hypoacetylation was specific to a particular coffee product, we first examined the effects of five commercially available coffee brands on H3K27ac levels. Immunoblot analysis demonstrated that all coffee brands significantly reduced H3K27ac levels in K562 cells, although the magnitude of the reduction varied among products (Fig. 3A). Similarly, both regular and decaffeinated coffee induced comparable decreases in H3K27ac (Fig. 3B), indicating that caffeine is not solely responsible for coffee-induced histone hypoacetylation. Because coffee treatment did not significantly alter the pH of the culture medium at the concentrations used (Fig. 3C), the observed reduction in histone acetylation is unlikely to result from nonspecific acidification of the culture environment.

**Figure 3.**
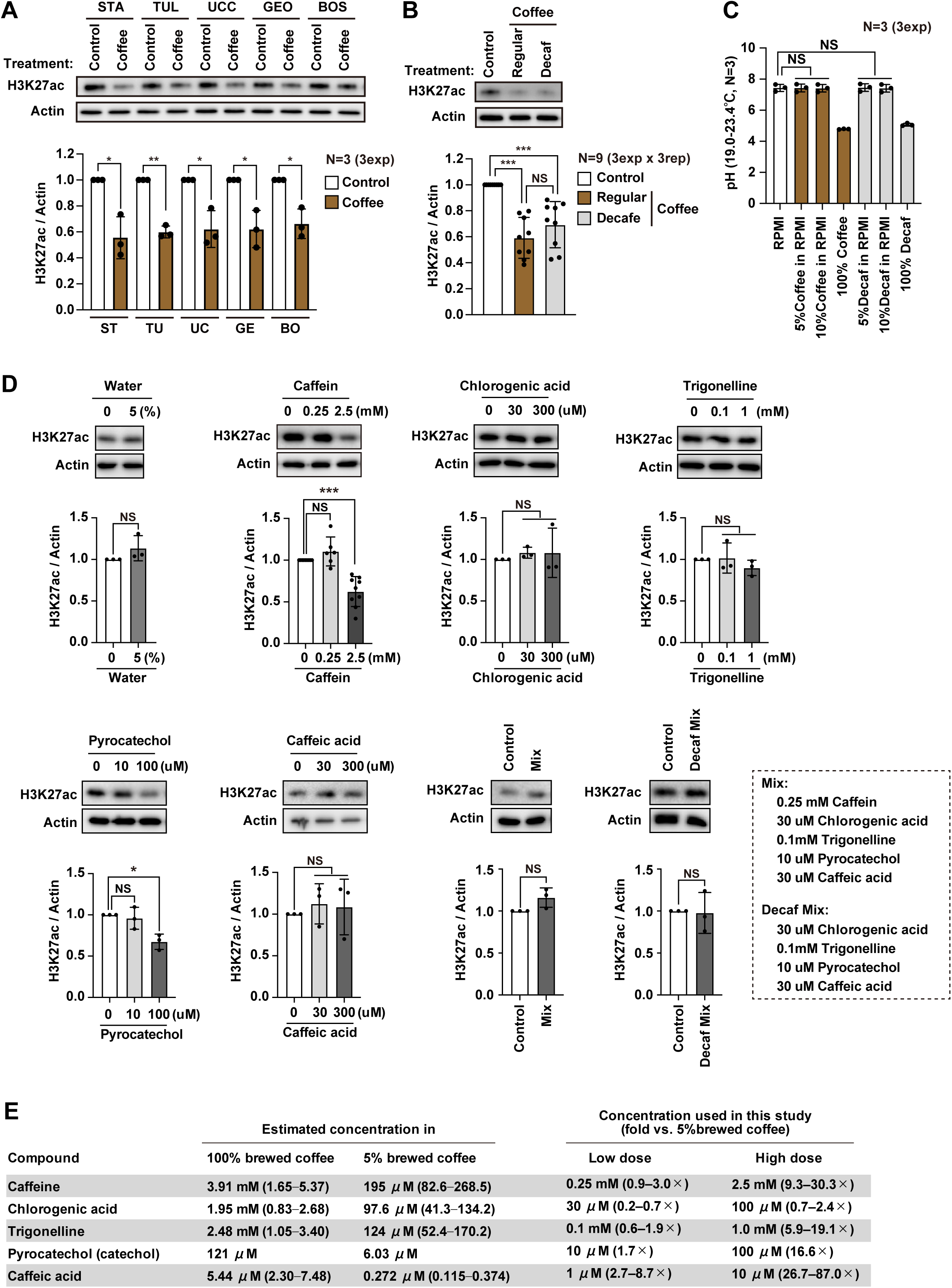
Identification of coffee-derived components contributing to histone hypoacetylation. **(A)** Immunoblot analysis of H3K27ac in K562 cells treated with five commercially available coffee brands (STA, Starbucks; TUL, Tully’s; UCC, UCC; GEO, Georgia; BOS, Boss). Representative immunoblots and corresponding quantification are shown. Data are presented as mean ± SD from three independent experiments. **(B)** Immunoblot analysis of H3K27ac in K562 cells treated with regular or decaffeinated coffee. Representative immunoblots and corresponding quantification are shown. Data are presented as mean ± SD from three independent experiments with three technical replicates each (n = 9 measurements). **(C)** pH of culture medium containing regular or decaffeinated coffee at the indicated concentrations. Data are presented as mean ± SD from three independent experiments. **(D)** Immunoblot analysis of H3K27ac in K562 cells treated with the indicated coffee-derived compounds or mixtures. Representative immunoblots and corresponding quantification are shown. Data are presented as mean ± SD from three independent experiments. **(E)** Estimated concentrations of major coffee-derived compounds in 100% brewed coffee and 5% brewed coffee, and the corresponding concentrations used for individual compound treatments in this study. Estimated concentrations were calculated from published measurements of brewed coffee, with the reported concentration ranges shown in parentheses. The low- and high-dose concentrations used for in vitro experiments are listed on the right.

We next investigated whether major coffee-derived compounds could reproduce the effects of whole coffee. Among the compounds examined, pyrocatechol and caffeine reduced H3K27ac levels only at relatively high concentrations, whereas chlorogenic acid, trigonelline, and caffeic acid had little or no detectable effect (Fig. 3D). Furthermore, a mixture containing chlorogenic acid (30 μM), trigonelline (0.1 mM), caffeic acid (30 μM), pyrocatechol (10 μM), and caffeine (0.25 mM) failed to fully reproduce the extent of histone hypoacetylation induced by whole coffee (Fig. 3D). Comparison of the experimental concentrations with the estimated concentrations in the 5% coffee preparation revealed that the tested concentrations ranged from approximately physiological levels to several-fold higher than those estimated in brewed coffee (Fig. 3E). These findings suggest that coffee-induced histone hypoacetylation cannot be explained by a single major constituent but is more likely mediated through the combined actions of multiple coffee-derived components, with pyrocatechol and caffeine representing potential contributing factors.

### Coffee induces histone hypoacetylation through coordinated regulation of histone acetyltransferases and histone deacetylases

To investigate the molecular mechanisms underlying coffee-induced histone hypoacetylation, we performed quantitative proteomic analysis of K562 cells following coffee treatment (Fig. 4A, B). Comparative proteomic profiling identified coordinated changes in proteins involved in histone acetylation regulation (Fig. 4C). Among histone acetyltransferases, quantitative proteomics showed reduced abundance of GCN5, which primarily acetylates H3K9 and H3K14, MOF, the major H4K16 acetyltransferase, and HAT1, which acetylates newly synthesized histone H4 at K5 and K12^15,49^, following coffee treatment (Fig. 4C). Immunoblot analysis confirmed significant reductions in GCN5 and MOF protein levels, whereas the reduction in HAT1 was not statistically significant (Fig. 4D).

**Figure 4.**
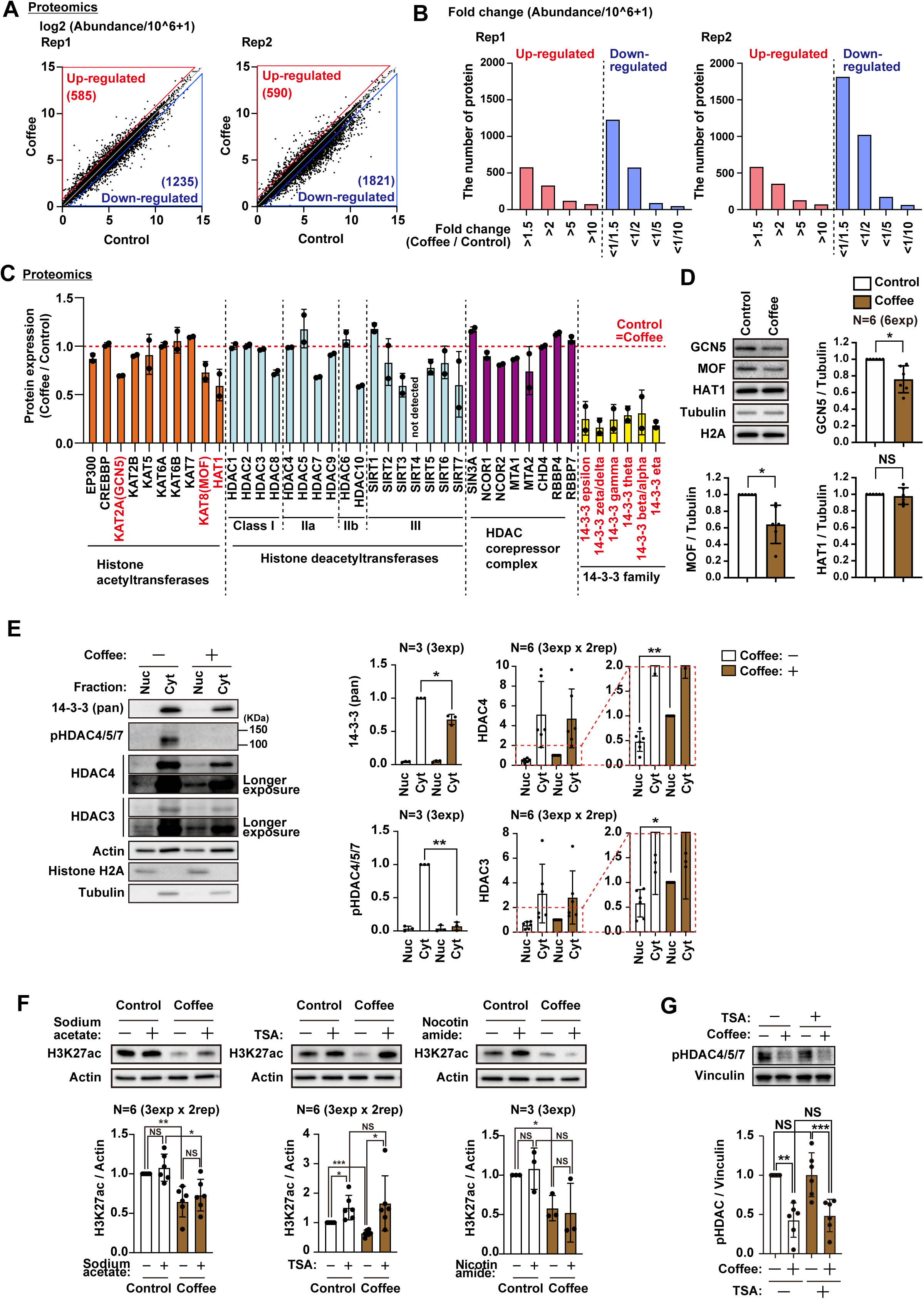
Molecular mechanisms underlying coffee-induced histone hypoacetylation. **(A)** Scatter plots showing global protein abundance determined by quantitative proteomic analysis in coffee-treated and control cells. Data from two independent biological replicates are shown. **(B)** Distribution of protein abundance changes in coffee-treated cells relative to controls for each biological replicate. **(C)** Relative protein expression of histone acetyltransferases, histone deacetylases, HDAC corepressor complex components, and 14-3-3 family proteins determined by quantitative proteomic analysis. Data are presented as mean ± SD from two biological replicates. **(D)** Immunoblot analysis of GCN5, MOF, and HAT1 in K562 cells following coffee treatment. Representative immunoblots and corresponding quantification are shown. Data are presented as mean ± SD from six independent experiments. **(E)** Subcellular fractionation followed by immunoblot analysis of 14-3-3 proteins, phosphorylated HDAC4/5/7 (pHDAC4/5/7), HDAC4, and HDAC3 in nuclear and cytoplasmic fractions. Representative immunoblots and corresponding quantification are shown. Data are presented as mean ± SD from the indicated numbers of independent experiments. **(F)** Immunoblot analysis showing the effects of 5 mM sodium acetate (24 h), 100 nM trichostatin A (TSA; 6 h), and 5 mM nicotinamide (24 h) on coffee-induced H3K27ac reduction in K562 cells. Representative immunoblots and corresponding quantification are shown. Data are presented as mean ± SD from six independent measurements (three independent experiments with two technical replicates each) for sodium acetate and TSA, and three independent experiments for nicotinamide.

Proteomic analysis further revealed reduced expression of multiple 14-3-3 family proteins (Fig. 4C). Because 14-3-3 proteins regulate the cytoplasmic retention of class IIa histone deacetylases through phosphorylation-dependent interactions^50–52^, we examined the intracellular localization of HDAC4, a representative class IIa HDAC, together with HDAC3, the catalytic component of the NCoR/SMRT corepressor complex recruited by class IIa HDACs ^16^. Coffee treatment reduced the abundance of 14-3-3 proteins and phosphorylated HDAC4/5/7 while increasing the nuclear accumulation of both HDAC4 and HDAC3 (Fig. 4E), suggesting that coffee promotes recruitment of the catalytically active HDAC3-containing corepressor complex to the nucleus, thereby enhancing histone deacetylation.

To determine whether histone deacetylases contribute functionally to coffee-induced histone hypoacetylation, cells were treated with the HDAC inhibitor trichostatin A (TSA), the class III HDAC inhibitor nicotinamide, or sodium acetate as a precursor for acetyl-CoA synthesis. TSA almost completely restored H3K27ac levels in coffee-treated cells, whereas neither nicotinamide nor sodium acetate produced a comparable effect (Fig. 4F). In contrast, TSA failed to restore the reduced phosphorylation of HDAC4/5/7 induced by coffee (Fig. 4G), indicating that expectedly decreased HDAC phosphorylation occurs upstream of histone hypoacetylation. Collectively, these findings suggest that coffee-induced histone hypoacetylation is primarily mediated by enhanced nuclear accumulation of histone deacetylases, while reduced expression of GCN5 and MOF may further contribute to this process.

### Promoter-associated active histone modifications are preferentially associated with transcriptional repression following coffee treatment

H3K27ac and H3K18ac are active histone acetylation marks enriched at active promoters and enhancers, whereas H3K4me3 marks active promoters and H3K36ac is associated with transcriptionally active gene bodies ^11,12^. Because coffee induced global reductions in these active histone modifications (Figs. 1 and 2), we next investigated whether genes marked by these modifications exhibit distinct transcriptional responses following coffee treatment. To address this question, genes were first grouped into four clusters according to promoter-associated H3K27ac, H3K18ac, and H3K4me3 signals. Gene body-associated H3K36ac and enhancer-associated H3K27ac and H3K18ac were analyzed in parallel (Fig. 5A). RNA-seq analysis revealed widespread transcriptional alterations that were highly reproducible across both independent replicates (Fig. 5B, C). Promoter-associated active histone modifications showed the strongest and most reproducible association with transcriptional repression across both biological replicates. Although gene body-associated H3K36ac showed a weaker association in one replicate, this effect was not consistently reproduced, whereas enhancer-associated active histone modifications showed little or no consistent association with transcriptional changes (Fig. 5D). Representative genome browser tracks demonstrated reduced promoter-associated H3K27ac together with decreased expression of *GALNT5, F2RL3, GBAP1, and CYB5RL* following coffee treatment (Fig. 5E, F). Collectively, these findings indicate that promoter-associated active histone modifications exhibit the strongest and most reproducible association with transcriptional repression following coffee treatment.

**Figure 5.**
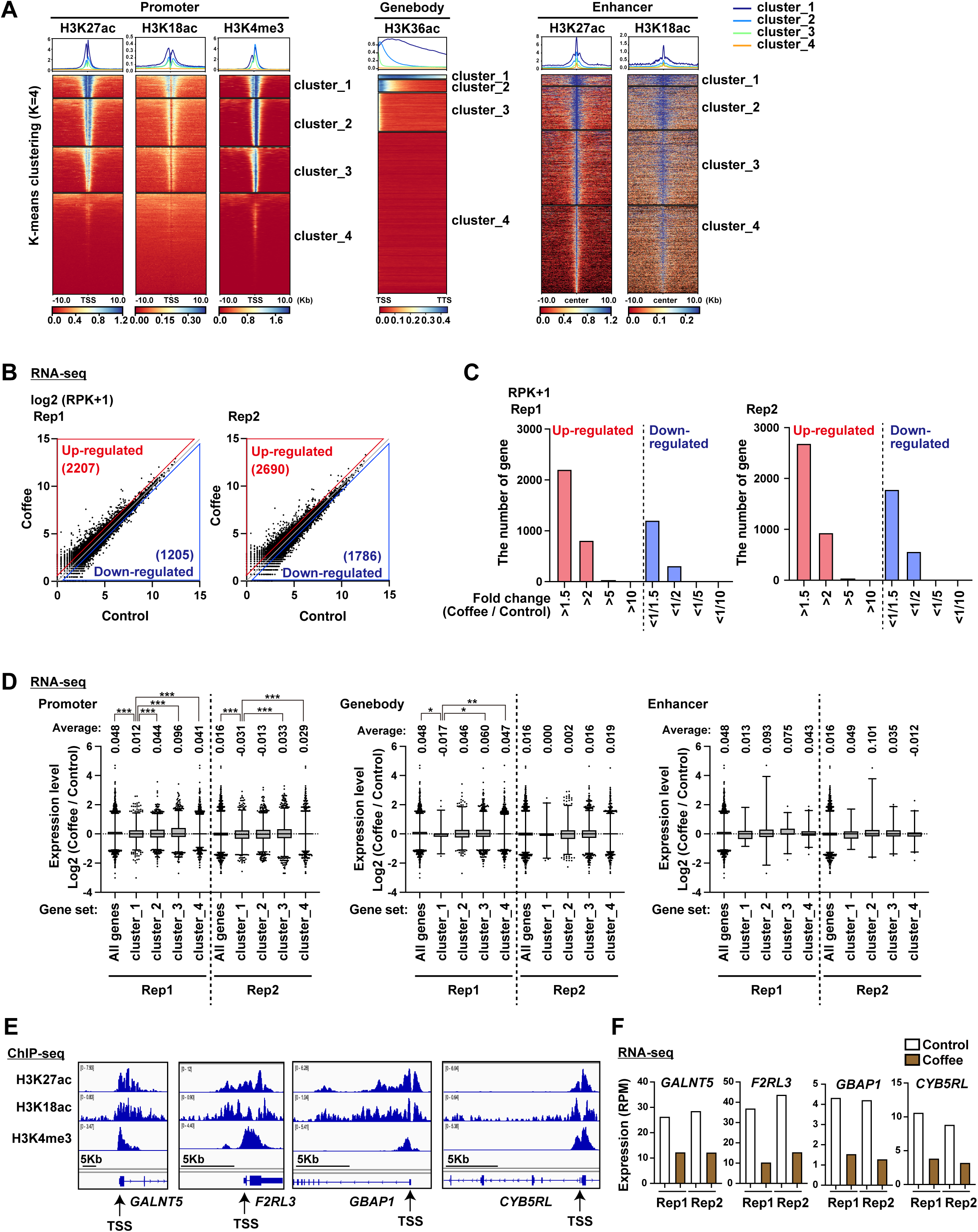
Genome-wide transcriptional alterations associated with histone modification-defined gene clusters following coffee treatment. **(A)** Heat maps and average profiles of promoter-associated H3K27ac, H3K18ac, and H3K4me3, gene body-associated H3K36ac, and enhancer-associated H3K27ac and H3K18ac. Genes were classified into four clusters (k = 4) based on histone modification enrichment profiles. **(B)** Scatter plots showing genome-wide gene expression determined by RNA-seq in control and coffee-treated K562 cells. Data from two independent biological replicates are shown. **(C)** Distribution of gene expression changes in coffee-treated cells relative to controls for each biological replicate. **(D)** RNA-seq expression changes of genes classified according to the histone modification-defined clusters shown in (A). Box plots show log₂(Coffee/Control) gene expression changes for all genes and for each cluster in two independent biological replicates. **(E)** Representative genome browser tracks showing H3K27ac, H3K18ac, and H3K4me3 ChIP-seq signals at the *GALNT5*, *F2RL3*, *GBAP1*, and *CYB5RL* loci. **(F)** RNA-seq expression levels of *GALNT5*, *F2RL3*, *GBAP1*, and *CYB5RL* in control and coffee-treated cells from two independent biological replicates.

### Coffee induces widespread loss of promoter-associated H3K27ac accompanied by transcriptional repression

Because promoter-associated active histone modifications exhibited the strongest association with transcriptional repression following coffee treatment (Fig. 5), we next focused on promoter-associated H3K27ac to identify genomic loci that lose this active histone mark. Consistent with the global reduction in H3K27ac observed by quantitative histone modification proteomics, immunoblotting, and immunofluorescence analyses (Figs. 1 and 2), chromatin immunoprecipitation (ChIP) demonstrated a marked reduction in the amount of H3K27ac-immunoprecipitated DNA following coffee treatment (Fig. 6A). ChIP-seq analysis further revealed a widespread decrease in H3K27ac signals surrounding transcription start sites in both independent replicates (Fig. 6B). To systematically identify promoters exhibiting reduced H3K27ac, we compared H3K27ac ChIP-seq CPM values within ±2 kb of transcription start sites between control and coffee-treated cells (Fig. 6C). Promoters with a control CPM value ≥ 0.3 and exhibiting greater than a twofold reduction following coffee treatment were defined as H3K27ac-lost promoters. Using these criteria, 4,785 and 4,755 H3K27ac-lost promoters were identified in Rep1 and Rep2, respectively, of which 4,263 were shared between both datasets (Fig. 6D). Gene Ontology analysis of the 4,263 shared H3K27ac-lost genes revealed significant enrichment of biological processes related to macromolecule metabolism, gene expression, RNA metabolism, translation, and cell cycle regulation (Fig. 6E). Integration of the ChIP-seq and RNA-seq datasets demonstrated that genes harboring H3K27ac-lost promoters exhibited significantly greater reductions in gene expression than genes without promoter H3K27ac loss in both independent replicates (Fig. 6F). Collectively, these findings demonstrate that coffee induces widespread loss of promoter-associated H3K27ac, providing a genome-wide framework for the transcriptional repression observed following coffee treatment.

**Figure 6.**
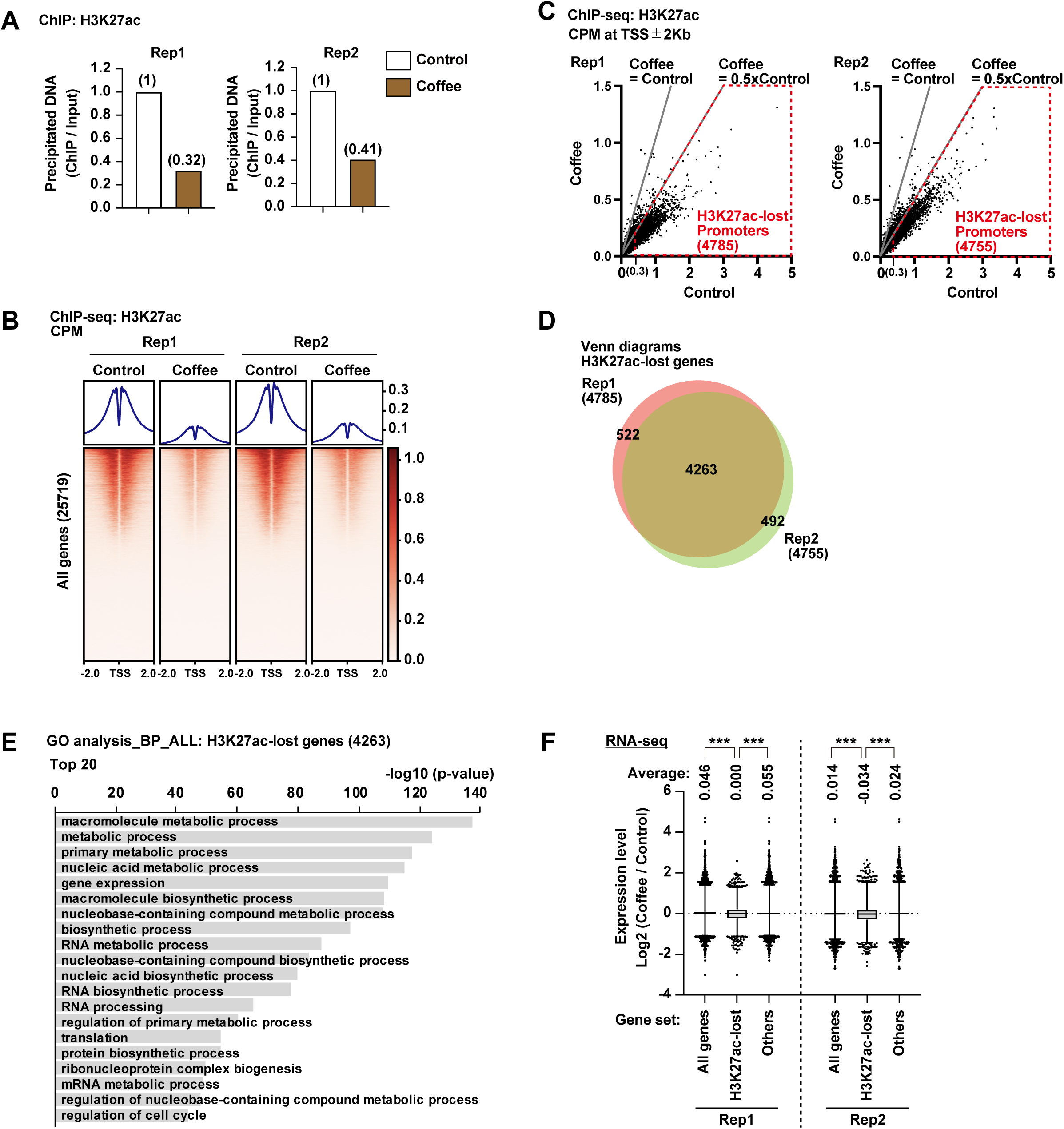
Genome-wide identification of promoter-associated H3K27ac loss and its transcriptional consequences following coffee treatment. **(A)** Relative amount of H3K27ac-immunoprecipitated DNA, reflecting global H3K27ac levels, in control and coffee-treated cells. Data from two independent biological replicates are shown. **(B)** Heat maps and average profiles of H3K27ac ChIP-seq signals across all annotated gene promoters (±2 kb from the transcription start site) in control and coffee-treated cells. **(C)** Scatter plots showing H3K27ac ChIP-seq CPM values at promoter regions (TSS ±2 kb) in control and coffee-treated cells. Genes with a promoter H3K27ac CPM ≥ 0.3 in control cells and showing a greater than twofold reduction in CPM following coffee treatment were defined as H3K27ac-lost promoters. **(D)** Venn diagram showing the overlap of H3K27ac-lost promoters identified in two independent biological replicates. **(E)** Gene Ontology (Biological Process) enrichment analysis of genes with H3K27ac-lost promoters shared between both biological replicates. **(F)** RNA-seq expression changes of genes with H3K27ac-lost promoters. Box plots show log₂(Coffee/Control) gene expression changes for all genes, H3K27ac-lost genes, and all remaining genes in two independent biological replicates.

### The MYC transcriptional program is preferentially affected by coffee-induced promoter H3K27ac loss

Having established that coffee induces widespread loss of promoter-associated H3K27ac accompanied by transcriptional repression (Fig. 6), we next sought to identify transcriptional programs preferentially affected by coffee treatment. Gene set enrichment analysis (GSEA) of both RNA-seq and quantitative proteomic datasets consistently identified MYC target gene sets (MYC Targets V1 and V2) as the most significantly negatively enriched pathways, whereas TNFα signaling via NF-κB, p53 pathway, and apoptosis were positively enriched (Fig. 7A). These findings suggested that suppression of the MYC transcriptional program represents a major molecular response to coffee treatment. Heatmap analysis further demonstrated coordinated downregulation of representative MYC target genes across both independent RNA-seq replicates (Fig. 7B). We therefore examined whether the *MYC* locus itself exhibited promoter H3K27ac loss. ChIP-seq genome browser visualization demonstrated a marked reduction in promoter-associated H3K27ac at the *MYC* locus in both independent replicates (Fig. 7C), which was further confirmed by ChIP analysis (Fig. 7D). Consistent with these chromatin changes, RNA-seq and quantitative proteomic analyses demonstrated reduced *MYC* expression following coffee treatment (Fig. 7E, F). RT-qPCR and immunoblot analyses further confirmed significant reductions in MYC mRNA and protein levels, respectively (Fig. 7G, H). We next examined representative MYC target genes that were consistently downregulated following coffee treatment. RT-qPCR confirmed significant reductions in the expression of *ODC1*, *LDHA*, *MCM7*, and *PRPS2* (Fig. 7I). Furthermore, ChIP-seq genome browser tracks together with ChIP analysis demonstrated marked reductions in promoter-associated H3K27ac at each of these loci (Fig. 7J, K). Collectively, these findings demonstrate that the MYC transcriptional program is preferentially affected by coffee-induced promoter H3K27ac loss, as evidenced by reduced promoter-associated H3K27ac at both the *MYC* locus and representative MYC target genes.

**Figure 7.**
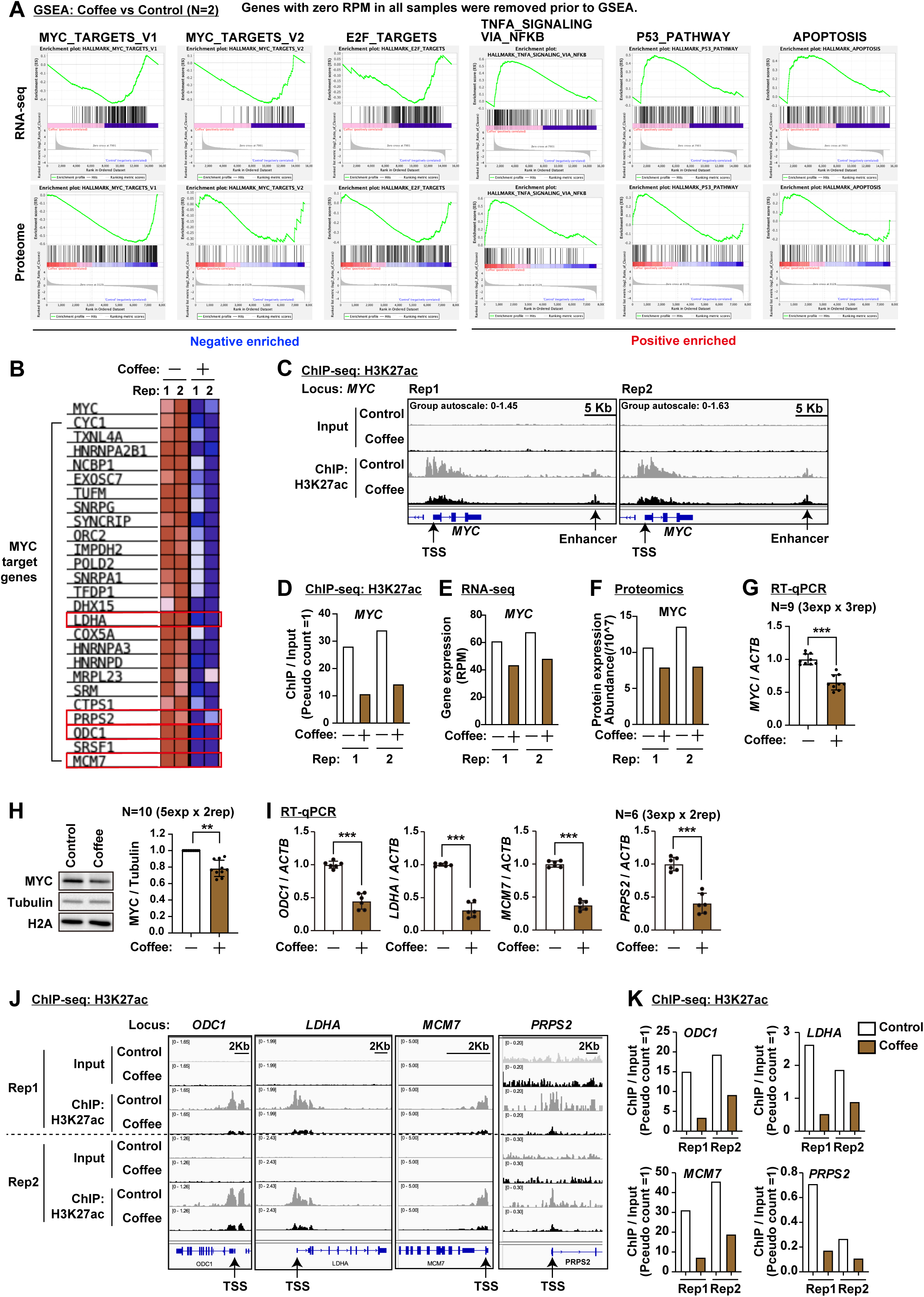
Suppression of the MYC transcriptional program is associated with reduced H3K27ac at MYC and MYC target loci following coffee treatment. **(A)** Gene Set Enrichment Analysis (GSEA) of RNA-seq and proteomic datasets showing enrichment of the indicated Hallmark gene sets. Genes with zero RPM across all RNA-seq samples were excluded prior to GSEA. **(B)** Heat map showing the expression of representative MYC target genes in control and coffee-treated cells determined by RNA-seq. **(C)** Representative genome browser tracks showing H3K27ac ChIP-seq signals at the *MYC* locus in two independent biological replicates. **(D)** H3K27ac ChIP-seq analysis at the *MYC* promoter. Data from two independent biological replicates are shown. **(E)** RNA-seq expression levels of *MYC* in control and coffee-treated cells from two independent biological replicates. **(F)** MYC protein abundance determined by quantitative proteomic analysis in control and coffee-treated cells from two independent biological replicates. **(G)** RT-qPCR analysis of *MYC* mRNA expression. Data are presented as mean ± SD from nine independent measurements (three independent experiments with three technical replicates each). **(H)** Immunoblot analysis of MYC protein in K562 cells following coffee treatment. Representative immunoblots and corresponding quantification are shown. Data are presented as mean ± SD from ten independent measurements (five independent experiments with two technical replicates each). **(I)** RT-qPCR analysis of representative MYC target genes (*ODC1, LDHA, MCM7,* and *PRPS2*). Data are presented as mean ± SD from six independent measurements (three independent experiments with two technical replicates each). **(J)** Representative genome browser tracks showing H3K27ac ChIP-seq signals at the *ODC1, LDHA, MCM7,* and *PRPS2* loci in two independent biological replicates. **(K)** H3K27ac ChIP analysis at the promoters of *ODC1, LDHA, MCM7,* and *PRPS2*. Data from two independent biological replicates are shown.

### Coffee suppresses cell proliferation independently of apoptosis

Having established that coffee induces global histone hypoacetylation and preferentially affects the MYC transcriptional program, we next investigated the cellular consequences of these molecular changes. Coffee treatment significantly reduced cell number and metabolic activity in K562 cells, as determined by cell counting and WST assays, respectively (Fig. 8A, B). Although cell death was modestly increased compared with untreated controls, the extent of cell death remained substantially lower than that induced by hydrogen peroxide, as determined by trypan blue exclusion assays (Fig. 8C, D). Consistent with these findings, coffee treatment did not induce cleavage of caspase-3, in contrast to hydrogen peroxide treatment, suggesting that the increased cell death induced by coffee is unlikely to be mediated by caspase-3-dependent apoptosis (Fig. 8E, F).

**Figure 8.**
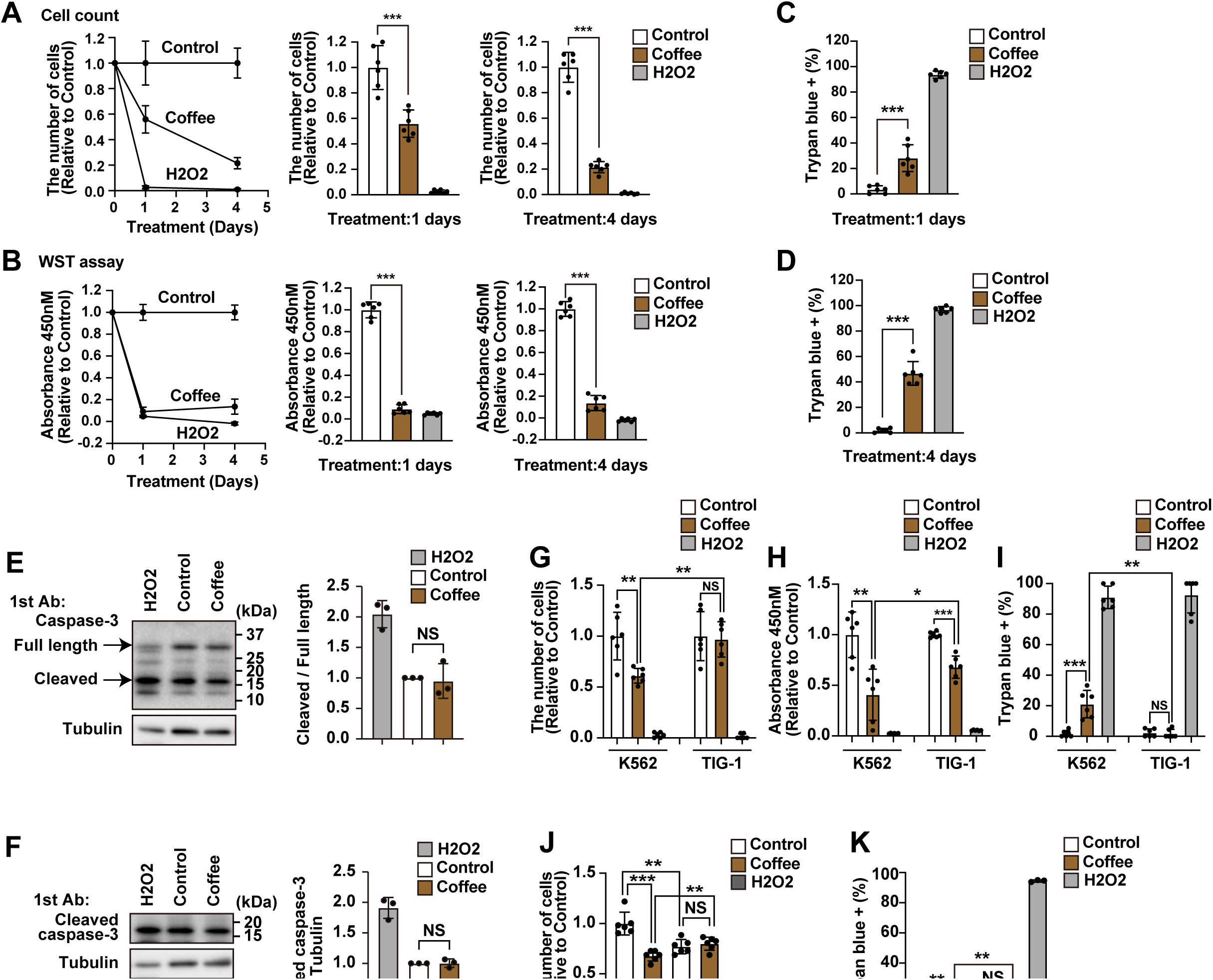
Coffee suppresses cell proliferation and viability independently of apoptosis. **(A)** Cell proliferation of K562 cells following coffee treatment, determined by direct cell counting. Time-course analysis and quantification after 1 and 4 days of treatment are shown. Data are presented as mean ± SD from the indicated independent experiments. **(B)** Cell viability determined by WST assay following coffee treatment. Time-course analysis and quantification after 1 and 4 days of treatment are shown. Data are presented as mean ± SD from the indicated independent experiments. **(C)** Percentage of trypan blue-positive K562 cells after 1 day of coffee treatment. Data are presented as mean ± SD from the indicated independent experiments. **(D)** Percentage of trypan blue-positive K562 cells after 4 days of coffee treatment. Data are presented as mean ± SD from the indicated independent experiments. **(E)** Immunoblot analysis of full-length and cleaved caspase-3 in K562 cells following coffee treatment. Representative immunoblots and corresponding quantification are shown. **(F)** Immunoblot analysis of cleaved caspase-3 in K562 cells using an antibody specific for cleaved caspase-3. Representative immunoblots and corresponding quantification are shown. **(G)** Cell proliferation of K562 and TIG-1 cells following coffee treatment, determined by direct cell counting. Data are presented as mean ± SD from the indicated independent experiments. **(H)** Cell viability of K562 and TIG-1 cells determined by WST assay following coffee treatment. Data are presented as mean ± SD from the indicated independent experiments. **(I)** Percentage of trypan blue-positive K562 and TIG-1 cells following coffee treatment. Data are presented as mean ± SD from the indicated independent experiments. **(J)** Effects of 100 nM trichostatin A (TSA) treatment for 6 h on coffee-induced inhibition of K562 cell proliferation. Data are presented as mean ± SD from the indicated independent experiments. **(K)** Effects of 100 nM trichostatin A (TSA) treatment for 6 h on the percentage of trypan blue-positive K562 cells following coffee treatment. Data are presented as mean ± SD from the indicated independent experiments.

**Figure 9.**
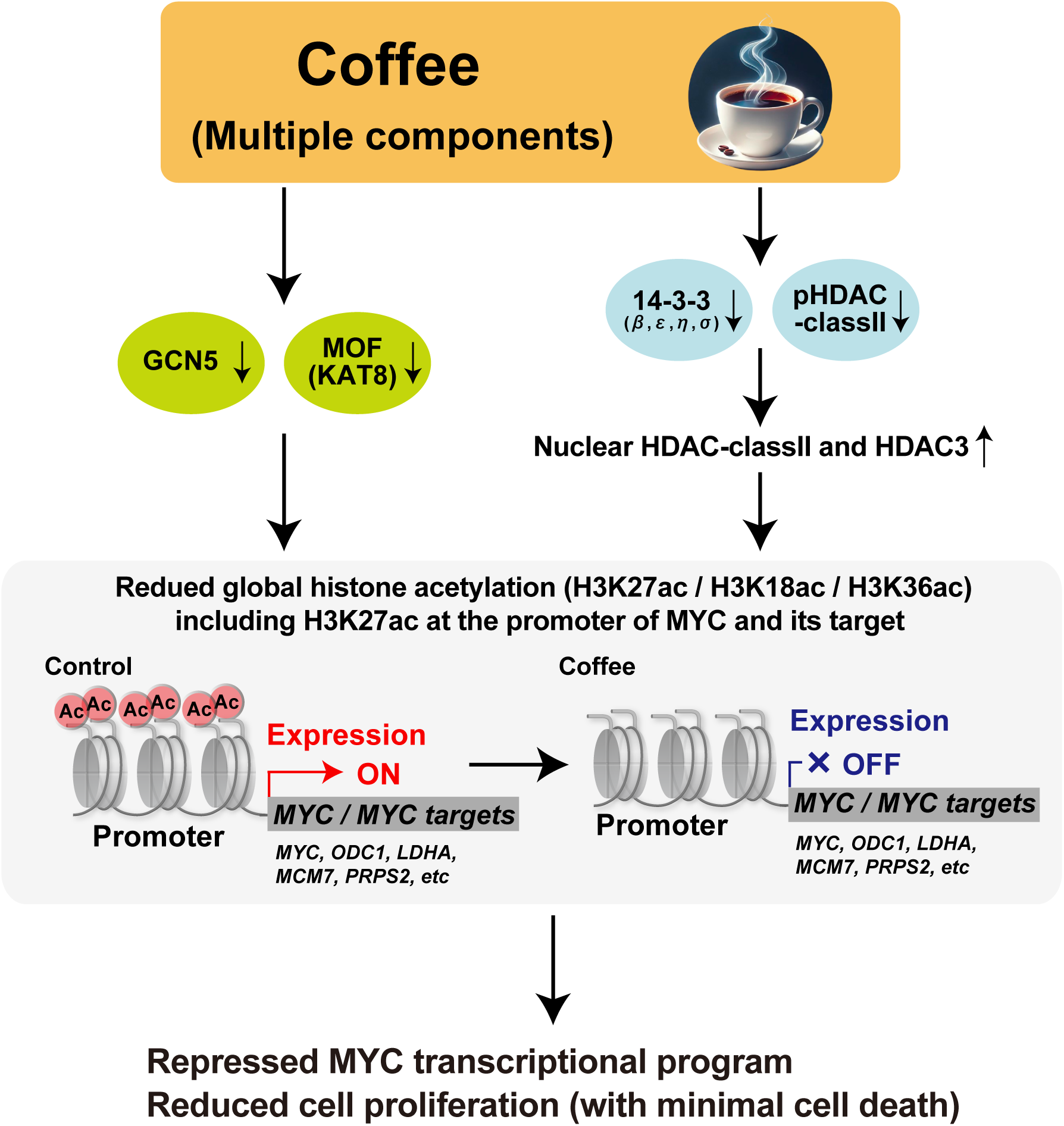
Proposed model of coffee-induced histone hypoacetylation and suppression of the MYC transcriptional program.

To determine whether these anti-proliferative effects are associated with histone hypoacetylation, we compared K562 cells, in which coffee induces marked histone hypoacetylation (Fig. 2A), with TIG-1 fibroblasts, in which histone acetylation remains unchanged following coffee treatment (Fig. 2B). Coffee treatment markedly reduced cell number and metabolic activity in K562 cells, whereas these effects were substantially weaker in TIG-1 cells (Fig. 8G, H). Consistent with this observation, coffee induced only a modest increase in cell death in K562 cells and no significant increase in TIG-1 cells (Fig. 8I). These findings suggest that the anti-proliferative effects of coffee are more pronounced in cells undergoing histone hypoacetylation, consistent with the suppression of the MYC transcriptional program observed in K562 cells.

Finally, because HDAC activity contributes to coffee-induced histone hypoacetylation (Fig. 4), we examined whether inhibition of HDAC activity influences the anti-proliferative effects of coffee. Treatment with trichostatin A (TSA) partially restored the reduction in cell number induced by coffee treatment without affecting cell death (Fig. 8J, K). Collectively, these findings demonstrate that coffee suppresses cell proliferation in association with histone hypoacetylation and suppression of the MYC transcriptional program, largely independently of caspase-3-dependent apoptosis. Furthermore, inhibition of HDAC activity partially attenuated the reduction in cell number induced by coffee, suggesting that HDAC activation contributes, at least in part, to the anti-proliferative effects of coffee.

## Discussion

In the present study, we demonstrate that coffee induces global histone hypoacetylation through coordinated alterations in histone acetyltransferases and histone deacetylases (Figs. 1–4). Multi-omics analyses integrating quantitative histone modification proteomics, quantitative proteomics, ChIP-seq, RNA-seq, and functional assays revealed that coffee induces genome-wide remodeling of active histone modifications, including H3K27ac, H3K18ac, H3K4me3, and H3K36ac (Figs. 1, 2, and 5). Genes marked by active promoter histone modifications, including H3K27ac, H3K18ac, and H3K4me3, exhibited greater transcriptional repression following coffee treatment (Fig. 5). Consistent with this finding, genes that lost promoter-associated H3K27ac were preferentially downregulated (Fig. 6), accompanied by preferential suppression of the MYC transcriptional program (Fig. 7) and reduced cell proliferation independent of detectable apoptosis (Fig. 8). Histone acetylation is a fundamental epigenetic mechanism regulating transcriptional activity and cellular identity^7,8,10^. Although numerous epidemiological and experimental studies have reported beneficial effects of coffee consumption on cancer, metabolic disorders, and age-related diseases ^3–6^, the underlying epigenetic mechanisms remain poorly understood. To the best of our knowledge, this is the first study to comprehensively demonstrate that whole coffee induces genome-wide histone hypoacetylation and to establish a molecular framework linking coordinated regulation of histone acetyltransferases and histone deacetylases with promoter H3K27ac loss, suppression of the MYC transcriptional program, and reduced cell proliferation.

Recent studies have shown that individual coffee-derived compounds can influence histone acetylation in specific biological contexts. Chronic caffeine intake was reported to alter genome-wide H3K27ac profiles in the mouse hippocampus in association with learning-dependent transcription^21^, whereas chlorogenic acid enhanced H3K27ac in human embryonic stem cells by promoting fatty acid β-oxidation and acetyl-CoA production.^22^. In contrast, our study demonstrates that whole coffee extract induces widespread histone hypoacetylation through coordinated regulation of both histone acetyltransferases and histone deacetylases. These differences may reflect the use of whole coffee rather than purified compounds, differences in cell type and physiological context, and distinct mechanisms of histone acetylation regulation. Indeed, although pyrocatechol and caffeine exhibited modest effects at higher concentrations, neither individual compounds nor a mixture of the major coffee-derived compounds reproduced the histone hypoacetylation induced by whole coffee (Fig. 3D). Comparison of the estimated concentrations of these compounds in the 5% coffee preparation with those used in our cell culture experiments further showed that the compounds were tested over a range extending from approximately 0.2- to 87-fold of their estimated concentrations in the 5% coffee preparation. (Fig. 3E). Therefore, the inability of these compounds, either individually or in combination, to reproduce the effects of whole coffee cannot simply be attributed to insufficient treatment concentrations. Rather, these findings argue against a dominant role for any single major constituent and suggest that the epigenetic activity of whole coffee is more likely mediated through cooperative interactions among multiple coffee-derived components and/or additional bioactive compounds generated during coffee roasting. These findings further emphasize that the biological effects of whole coffee cannot be adequately explained by isolated constituents alone, highlighting the importance of evaluating whole-food preparations when investigating the epigenetic effects of dietary factors.

Recent studies have begun to identify direct molecular targets of coffee-derived compounds. For example, caffeic acid, a major metabolite of chlorogenic acid, was recently shown to directly bind ribosomal protein S5 (RPS5), suppress cyclin D1 expression, and inhibit colorectal cancer cell growth^53^. Collectively, these findings suggest that previous studies have primarily identified individual molecular targets or biological activities of specific coffee-derived compounds, whereas our study reveals that whole coffee exerts higher-order regulation of the epigenetic landscape through coordinated remodeling of histone acetylation. This global chromatin remodeling was accompanied by widespread transcriptional reprogramming, including suppression of the MYC transcriptional program.

Our findings suggest that coffee-induced histone hypoacetylation is mediated by coordinated regulation of both histone acetyltransferases and histone deacetylases (Fig. 4). Quantitative proteomic and immunoblot analyses consistently demonstrated reduced expression of the histone acetyltransferases GCN5 and MOF (Fig. 4C, D), whereas coffee also promoted nuclear accumulation of class IIa HDACs accompanied by reduced phosphorylation of HDAC4/5/7 and decreased abundance of 14-3-3 proteins (Fig. 4C, E). Furthermore, pharmacological inhibition of HDACs with trichostatin A (TSA) restored histone acetylation without rescuing HDAC phosphorylation (Fig. 4F, G), suggesting that HDAC dephosphorylation occurs upstream of histone hypoacetylation. Interestingly, the reduction in phosphorylated HDAC4/5/7 appeared more pronounced than the decrease in 14-3-3 proteins (Fig. 4E). Because phosphorylation of class IIa HDACs is required for their interaction with 14-3-3 proteins and cytoplasmic retention^50–52^, these observations raise the possibility that HDAC dephosphorylation precedes the reduction of 14-3-3 proteins. Loss of HDAC phosphorylation may disrupt HDAC–14-3-3 interactions, leading to nuclear accumulation of HDACs and subsequent destabilization of unbound 14-3-3 proteins. Although this model remains speculative, it is consistent with our observation that TSA restored histone acetylation without affecting HDAC phosphorylation (Fig. 4F). More broadly, these findings raise the possibility that coffee influences cellular phosphorylation signaling beyond HDAC regulation. Future phosphoproteomic analyses, together with genetic and chemical screening approaches, will be important for identifying the upstream kinases, phosphatases, and coffee-derived compounds responsible for coffee-induced histone hypoacetylation.

Another notable finding of this study is that widespread histone hypoacetylation was not accompanied by a corresponding global increase in histone methylation (Fig. 1). This suggests that coffee primarily attenuates active chromatin rather than converting it into a fully repressive chromatin state. Consistent with this interpretation, RNA-seq revealed predominantly moderate reductions in gene expression rather than complete transcriptional silencing. These findings suggest that coffee primarily attenuates active chromatin without converting it into a globally repressive chromatin state. Such epigenetic changes may represent a relatively reversible and plastic mode of chromatin regulation.

An important finding of this study is that promoter-associated active histone modifications exhibited a substantially stronger association with transcriptional repression than enhancer-associated histone acetylation (Figs. 5 and 6). Among the active histone modifications examined, promoter-associated H3K27ac showed the closest association with reduced gene expression and therefore became the focus of our subsequent analyses. This preferential association was also evident at the MYC locus, where promoter-associated H3K27ac was markedly reduced whereas enhancer-associated H3K27ac was largely preserved (Fig. 7). Consistent with the established role of promoter-associated histone modifications in transcriptional regulation, these findings indicate that coffee-induced changes in promoter-associated H3K27ac are more closely linked to gene expression than corresponding changes at enhancers. Although the molecular basis for this preference remains unclear, preferential remodeling of promoter-associated histone modifications may contribute to the transcriptional effects of coffee. Notably, this epigenetic response was not universal across cell types, as coffee-induced histone hypoacetylation was observed in K562 cells but not in TIG-1 fibroblasts or HCT116 cells, suggesting that the response depends on cellular context rather than reflecting a nonspecific effect of coffee treatment.

Among the transcriptional programs affected by coffee, the MYC transcriptional program was the most prominently suppressed pathway identified by both transcriptomic and proteomic analyses (Fig. 7C). MYC is a master transcription factor that regulates cell proliferation, metabolism, ribosome biogenesis, and cell-cycle progression, and dysregulated MYC activity is a hallmark of many human cancers ^54,55^. In the present study, coffee was associated not only with reduced *MYC* expression but also with decreased promoter-associated H3K27ac at both the *MYC* locus and representative MYC target genes. These observations suggest that remodeling of promoter-associated histone acetylation may contribute to coordinated suppression of the MYC transcriptional program. Given that aberrant MYC activation has also been implicated in aging and age-associated diseases ^56^, the epigenetic regulation of MYC identified in this study may provide a mechanistic link between coffee consumption and its reported beneficial effects on cancer and healthy aging. More broadly, accumulating evidence indicates that age-associated changes in chromatin organization and histone modifications are fundamental features of the aging process and contribute to the regulation of stress responses and tissue homeostasis ^57,58^. In this context, our finding that coffee remodels the histone acetylation landscape raises the possibility that coffee may act as a dietary epigenetic modulator, influencing biological processes beyond MYC regulation. Although the physiological relevance of these epigenetic changes remains to be established in vivo, modulation of histone acetylation by coffee may represent one mechanism through which coffee contributes to healthy aging and other health benefits reported in epidemiological studies.

Several limitations of this study should be acknowledged. First, our findings were obtained primarily using cultured cell lines and therefore require validation in vivo. Second, although pyrocatechol and caffeine partially reproduced histone hypoacetylation at relatively high concentrations (Fig. 3D), the coffee-derived components responsible for the full biological activity remain to be identified. Future studies integrating phosphoproteomics, genome-wide CRISPR screening, and chemical screening approaches will be valuable for identifying the upstream signaling pathways and coffee-derived molecules responsible for coffee-induced histone hypoacetylation.

In conclusion, our study identifies global remodeling of active histone modifications, characterized predominantly by histone hypoacetylation, as a previously unrecognized epigenetic response to coffee and establishes a molecular framework linking histone acetylation remodeling with suppression of the MYC transcriptional program and reduced cell proliferation. These findings provide a foundation for understanding the epigenetic mechanisms underlying the biological effects of coffee and offer new perspectives for investigating coffee-derived compounds as modulators of chromatin regulation.

## Acknowledgments

We thank Dr. Noritaka Yamaguchi (Meiji Pharmaceutical University) for kindly providing the MCF-7, A549, A431, and HCT116 cell lines used in this study. The authors thank Dr. Megumi Tago and Dr. Yosuke Nakazawa (Keio University) for kindly providing the K562 cell line and for general support. The authors thank Junko Yamada for maintaining the shared equipment platform and for her logistical support. The authors thank Atsushi Takeda, Junko Takeda, Shoji Aoyama, Hiromu Iwaori, Eri Iwaori, and Yumi Aoyama for their valuable discussions and insightful comments from a consumer perspective, which contributed to the conception and experimental design of this study. This work was supported in part by JSPS KAKENHI (Grant Number 23K07851); the Kobayashi Foundation; the Friends of Leukemia Research Fund (Takaku Fumimaro Award); the Japanese Society of Hematology; the Keio University Academic Development Fund; the Keio University Fukuzawa Fund; the Chemo-Sero-Therapeutic Research Institute (Kaketsuken); the Suzuken Memorial Foundation; the Mochida Memorial Foundation; and the Takeda Science Foundation. ChatGPT (OpenAI) was used under author supervision for minor revisions, consistency checking, and idea exploration. All content was verified by the authors.

## Authorship Contributions

Y.C. and K.A. contributed equally to this work. Y.C. and K.A. performed the experiments, analyzed the data, and prepared the figures. M.M.-Y., Y.N.-T., T.Y., M.O., and A.I. contributed to and supported the ChIP-seq experiments and provided valuable scientific advice and critical comments on the manuscript. K.A. conceived and supervised the study, secured funding, analyzed the data, and wrote the manuscript. All authors reviewed and approved the final manuscript.

## References

1 Bunge, A. C., Mazac, R., Clark, M. & Gordon, L. Emerging alternatives to coffee, cocoa and palm oil deserve a spot on the research agenda. Nat Food 6, 2–5 (2025). 10.1038/s43016-024-01103-w

2 Salojärvi, J. et al. The genome and population genomics of allopolyploid Coffea arabica reveal the diversification history of modern coffee cultivars. Nat Genet 56, 721–731 (2024). 10.1038/s41588-024-01695-w

3 Poole, R. et al. Coffee consumption and health: umbrella review of meta-analyses of multiple health outcomes. Bmj 359, j5024 (2017). 10.1136/bmj.j5024

4 van Dam, R. M., Hu, F. B. & Willett, W. C. Coffee, Caffeine, and Health. The New England journal of medicine 383, 369–378 (2020). 10.1056/NEJMra1816604

5 Ludwig, I. A., Clifford, M. N., Lean, M. E., Ashihara, H. & Crozier, A. Coffee: biochemistry and potential impact on health. Food Funct 5, 1695–1717 (2014). 10.1039/c4fo00042k

6 Nieber, K. The Impact of Coffee on Health. Planta Med 83, 1256–1263 (2017). 10.1055/s-0043-115007

7 Allis, C. D. & Jenuwein, T. The molecular hallmarks of epigenetic control. Nat Rev Genet 17, 487–500 (2016). 10.1038/nrg.2016.59

8 Berger, S. L. The complex language of chromatin regulation during transcription. Nature 447, 407–412 (2007). 10.1038/nature05915

9 Jaenisch, R. & Bird, A. Epigenetic regulation of gene expression: how the genome integrates intrinsic and environmental signals. Nat Genet 33 **Suppl**, 245–254 (2003). 10.1038/ng1089

10 Kouzarides, T. Chromatin modifications and their function. Cell 128, 693–705 (2007). 10.1016/j.cell.2007.02.005

11 Creyghton, M. P. et al. Histone H3K27ac separates active from poised enhancers and predicts developmental state. Proceedings of the National Academy of Sciences of the United States of America 107, 21931–21936 (2010). 10.1073/pnas.1016071107

12 Kundaje, A. et al. Integrative analysis of 111 reference human epigenomes. Nature 518, 317–330 (2015). 10.1038/nature14248

13 de Ruijter, A. J., van Gennip, A. H., Caron, H. N., Kemp, S. & van Kuilenburg, A. B. Histone deacetylases (HDACs): characterization of the classical HDAC family. Biochem J 370, 737–749 (2003). 10.1042/bj20021321

14 Frank, S. R. et al. MYC recruits the TIP60 histone acetyltransferase complex to chromatin. EMBO Rep 4, 575–580 (2003). 10.1038/sj.embor.embor861

15 Roth, S. Y., Denu, J. M. & Allis, C. D. Histone acetyltransferases. Annual review of biochemistry 70, 81–120 (2001). 10.1146/annurev.biochem.70.1.81

16 Haberland, M., Montgomery, R. L. & Olson, E. N. The many roles of histone deacetylases in development and physiology: implications for disease and therapy. Nat Rev Genet 10, 32–42 (2009). 10.1038/nrg2485

17 Choi, S. W. & Friso, S. Epigenetics: A New Bridge between Nutrition and Health. Adv Nutr 1, 8–16 (2010). 10.3945/an.110.1004

18 Chuang, Y. H. et al. Coffee consumption is associated with DNA methylation levels of human blood. Eur J Hum Genet 25, 608–616 (2017). 10.1038/ejhg.2016.175

19 Karabegović, I. et al. Epigenome-wide association meta-analysis of DNA methylation with coffee and tea consumption. Nat Commun 12, 2830 (2021). 10.1038/s41467-021-22752-6

20 Ding, Q., Xu, Y. M. & Lau, A. T. Y. The Epigenetic Effects of Coffee. Molecules 28 (2023). 10.3390/molecules28041770

21 Paiva, I. et al. Caffeine intake exerts dual genome-wide effects on hippocampal metabolism and learning-dependent transcription. J Clin Invest 132 (2022). 10.1172/jci149371

22 Zong, M. et al. Chlorogenic acid promotes fatty acid beta-oxidation to increase hESCs proliferation and lipid synthesis. Scientific reports 15, 7095 (2025). 10.1038/s41598-025-91582-z

23 Funakoshi-Tago, M. et al. Pyrocatechol, a component of coffee, suppresses LPS-induced inflammatory responses by inhibiting NF-κB and activating Nrf2. Scientific reports 10, 2584 (2020). 10.1038/s41598-020-59380-x

24 Murata, T. et al. Suppression of Neuroinflammation by Coffee Component Pyrocatechol via Inhibition of NF-κB in Microglia. International journal of molecular sciences 25 (2023). 10.3390/ijms25010316

25 Nakayama, T., Funakoshi-Tago, M. & Tamura, H. Coffee reduces KRAS expression in Caco-2 human colon carcinoma cells via regulation of miRNAs. Oncol Lett 14, 1109–1114 (2017). 10.3892/ol.2017.6227

26 Fujioka, K. & Shibamoto, T. Chlorogenic acid and caffeine contents in various commercial brewed coffees. Food Chemistry 106, 217–221 (2008). 10.1016/j.foodchem.2007.05.091

27 Rodrigues, N. P. & Bragagnolo, N. Identification and quantification of bioactive compounds in coffee brews by HPLC–DAD–MSn. Journal of Food Composition and Analysis 32, 105–115 (2013). 10.1016/j.jfca.2013.09.002

28 Kaito, S. et al. Inhibition of TOPORS ubiquitin ligase augments the efficacy of DNA hypomethylating agents through DNMT1 stabilization. Nat Commun 15, 7359 (2024). 10.1038/s41467-024-50498-4

29 Yuki, R. et al. Desuppression of TGF-β signaling via nuclear c-Abl-mediated phosphorylation of TIF1γ/TRIM33 at Tyr-524, -610, and -1048. Oncogene 38, 637–655 (2019). 10.1038/s41388-018-0481-z

30 Mashimo, M. et al. PARP1 inhibition alleviates injury in ARH3-deficient mice and human cells. JCI insight 4 (2019). 10.1172/jci.insight.124519

31 Aoyama, K. et al. c-Abl induces stabilization of histone deacetylase 1 (HDAC1) in a kinase activity-dependent manner. Cell biology international 39, 446–456 (2015). 10.1002/cbin.10413

32 Honda, T. et al. Protective role for lipid modifications of Src-family kinases against chromosome missegregation. Scientific reports 6 (2016). 10.1038/srep38751

33 Schneider, C. A., Rasband, W. S. & Eliceiri, K. W. NIH Image to ImageJ: 25 years of image analysis. Nat Methods 9, 671–675 (2012). 10.1038/nmeth.2089

34 Kubota, S. et al. Role for Tyrosine Phosphorylation of A-kinase Anchoring Protein 8 (AKAP8) in Its Dissociation from Chromatin and the Nuclear Matrix. The Journal of biological chemistry 290, 10891–10904 (2015). 10.1074/jbc.M115.643882

35 Aoyama, K. et al. Formation of long and winding nuclear F-actin bundles by nuclear c-Abl tyrosine kinase. Experimental cell research 319, 3251–3268 (2013). 10.1016/j.yexcr.2013.09.003

36 Higashi, K., Tanaka, Y., Kosako, H. & Aoyama, K. Identification of MRVI1-Interacting Proteins by Biotin-Based Proximity Labeling Reveals NPM-ALK-Dependent Interaction Dynamics. J Biochem (2025). 10.1093/jb/mvaf057

37 Aoyama, K. et al. Nuclear c-Abl-mediated tyrosine phosphorylation induces chromatin structural changes through histone modifications that include H4K16 hypoacetylation. Experimental cell research 317, 2874–2903 (2011). 10.1016/j.yexcr.2011.09.013

38 Kubota, S. et al. Phosphorylation of KRAB-associated protein 1 (KAP1) at Tyr-449, Tyr-458, and Tyr-517 by nuclear tyrosine kinases inhibits the association of KAP1 and heterochromatin protein 1α (HP1α) with heterochromatin. The Journal of biological chemistry 288, 17871–17883 (2013). 10.1074/jbc.M112.437756

39 Aoyama, K. et al. Ezh1 Targets Bivalent Genes to Maintain Self-Renewing Stem Cells in Ezh2-Insufficient Myelodysplastic Syndrome. iScience 9, 161–174 (2018). 10.1016/j.isci.2018.10.008

40 Aoyama, K. et al. PRC2 insufficiency causes p53-dependent dyserythropoiesis in myelodysplastic syndrome. Leukemia 35, 1156–1165 (2021). 10.1038/s41375-020-01023-1

41 Mochizuki-Kashio, M. et al. Ezh2 loss in hematopoietic stem cells predisposes mice to develop heterogeneous malignancies in an Ezh1-dependent manner. Blood 126, 1172–1183 (2015). 10.1182/blood-2015-03-634428

42 Zou, Z., Ohta, T. & Oki, S. ChIP-Atlas 3.0: a data-mining suite to explore chromosome architecture together with large-scale regulome data. Nucleic Acids Res 52, W45–w53 (2024). 10.1093/nar/gkae358

43 Tanaka, T. et al. Internal deletion of BCOR reveals a tumor suppressor function for BCOR in T lymphocyte malignancies. The Journal of experimental medicine 214, 2901–2913 (2017). 10.1084/jem.20170167

44 Castanza, A. S. et al. Extending support for mouse data in the Molecular Signatures Database (MSigDB). Nat Methods 20, 1619–1620 (2023). 10.1038/s41592-023-02014-7

45 Huang da, W., Sherman, B. T. & Lempicki, R. A. Systematic and integrative analysis of large gene lists using DAVID bioinformatics resources. Nat Protoc 4, 44–57 (2009). 10.1038/nprot.2008.211

46 Huang, D. W. et al. DAVID Bioinformatics Resources: expanded annotation database and novel algorithms to better extract biology from large gene lists. Nucleic Acids Res 35, W169–175 (2007). 10.1093/nar/gkm415

47 Higashi, K., Cho, Y. & Aoyama, K. Multifaceted MRVI1 Serves as a Tumor Suppressor in HCT116 Colorectal Cancer Cells. Biol Pharm Bull 49, 589–593 (2026). 10.1248/bpb.b25-00756

48 Yuki, R. et al. Overexpression of zinc-finger protein 777 (ZNF777) inhibits proliferation at low cell density through down-regulation of FAM129A. J Cell Biochem 116, 954–968 (2015). 10.1002/jcb.25046

49 Marmorstein, R. & Zhou, M. M. Writers and readers of histone acetylation: structure, mechanism, and inhibition. Cold Spring Harbor perspectives in biology 6, a018762 (2014). 10.1101/cshperspect.a018762

50 Grozinger, C. M. & Schreiber, S. L. Regulation of histone deacetylase 4 and 5 and transcriptional activity by 14-3-3-dependent cellular localization. Proceedings of the National Academy of Sciences of the United States of America 97, 7835–7840 (2000). 10.1073/pnas.140199597

51 McKinsey, T. A., Zhang, C. L., Lu, J. & Olson, E. N. Signal-dependent nuclear export of a histone deacetylase regulates muscle differentiation. Nature 408, 106–111 (2000). 10.1038/35040593

52 Parra, M. Class IIa HDACs - new insights into their functions in physiology and pathology. Febs j 282, 1736–1744 (2015). 10.1111/febs.13061

53 Watanabe, M. et al. Caffeic acid suppresses cyclin D1 expression by directly binding to ribosomal protein S5 in colorectal cancer cells. Scientific reports 16 (2026). 10.1038/s41598-026-42196-6

54 Dang, C. V. MYC on the path to cancer. Cell 149, 22–35 (2012). 10.1016/j.cell.2012.03.003

55 Dhanasekaran, R. et al. The MYC oncogene - the grand orchestrator of cancer growth and immune evasion. Nat Rev Clin Oncol 19, 23–36 (2022). 10.1038/s41571-021-00549-2

56 Hofmann, J. W. et al. Reduced expression of MYC increases longevity and enhances healthspan. Cell 160, 477–488 (2015). 10.1016/j.cell.2014.12.016

57 Aoyama, K., Itokawa, N., Oshima, M. & Iwama, A. Epigenetic Memories in Hematopoietic Stem and Progenitor Cells. Cells 11 (2022). 10.3390/cells11142187

58 Itokawa, N. et al. Epigenetic traits inscribed in chromatin accessibility in aged hematopoietic stem cells. Nat Commun 13, 2691 (2022). 10.1038/s41467-022-30440-2

