## Supplementary Information for "Coffee remodels the global histone acetylation landscape with reduced H3K27ac at the *MYC* promoter"

| Compound | Fujioka et al. (mg/g ground coffee) | Rodrigues et al. (mg/100 g dry extract) | Ratio to total CGA | Estimated (mg/g ground coffee) | MW (g/mol) | Estimated concentration in 100% coffee extract (mM) | Range (mM) | Estimated concentration in 5% coffee extract (μM) | Range (μM) |
|---|---|---|---|---|---|---|---|---|---|
| Total chlorogenic acids | 5.26–17.1 (12.45) | 4162 | 1 | 12.45 | 354.31 | 1.952 | 0.825–2.684 | 97.6 | 41.3–134.2 |
| Chlorogenic acid lactones | — | 779 | 0.18717 | 2.33 | 336.3 | 0.385 | 0.163–0.529 | 19.3 | 8.15–26.5 |
| *p*-Coumaric acid | — | 2.4 | 0.000577 | 0.00718 | 164.16 | 0.00243 | 0.00103–0.00334 | 0.121 | 0.0515–0.167 |
| Trigonelline | — | 2044 | 0.49111 | 6.114 | 137.14 | 2.477 | 1.047–3.404 | 123.9 | 52.4–170.2 |
| Nicotinic acid | — | 100.4 | 0.024123 | 0.3 | 123.11 | 0.1355 | 0.0573–0.1863 | 6.78 | 2.87–9.32 |
| Caffeine | — | 4565 | 1.096828 | 13.656 | 194.19 | 3.907 | 1.652–5.370 | 195.3 | 82.6–268.5 |
| Theobromine | — | 12.5 | 0.003003 | 0.03739 | 180.16 | 0.01153 | 0.00487–0.01584 | 0.577 | 0.244–0.792 |
| Caffeic acid | — | 5.9 | 0.001418 | 0.01765 | 180.16 | 0.00544 | 0.00230–0.00748 | 0.272 | 0.115–0.374 |
| Pyrocatechol † | — | — | — | — | 110.11 | 0.1206 | — | 6.03 | — |
